## Supplement Figures 1, 3 & 4 for "Subchronic alteration of vestibular hair cells in mice: implications for multisensory gaze stabilization"

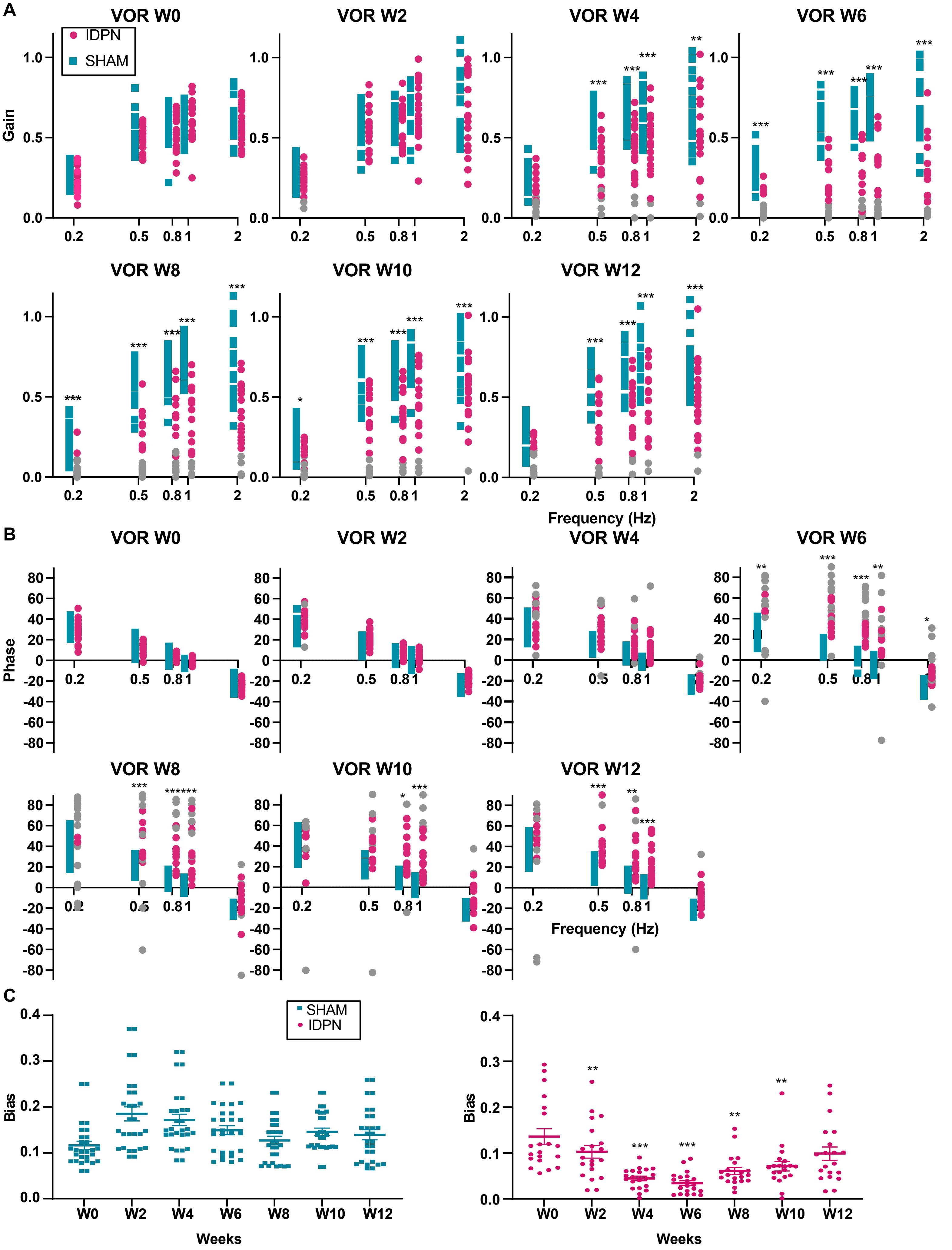


**Figure 1 supplement: Effects of subchronic IDPN on canalar- and otolith-dependent VOR.**

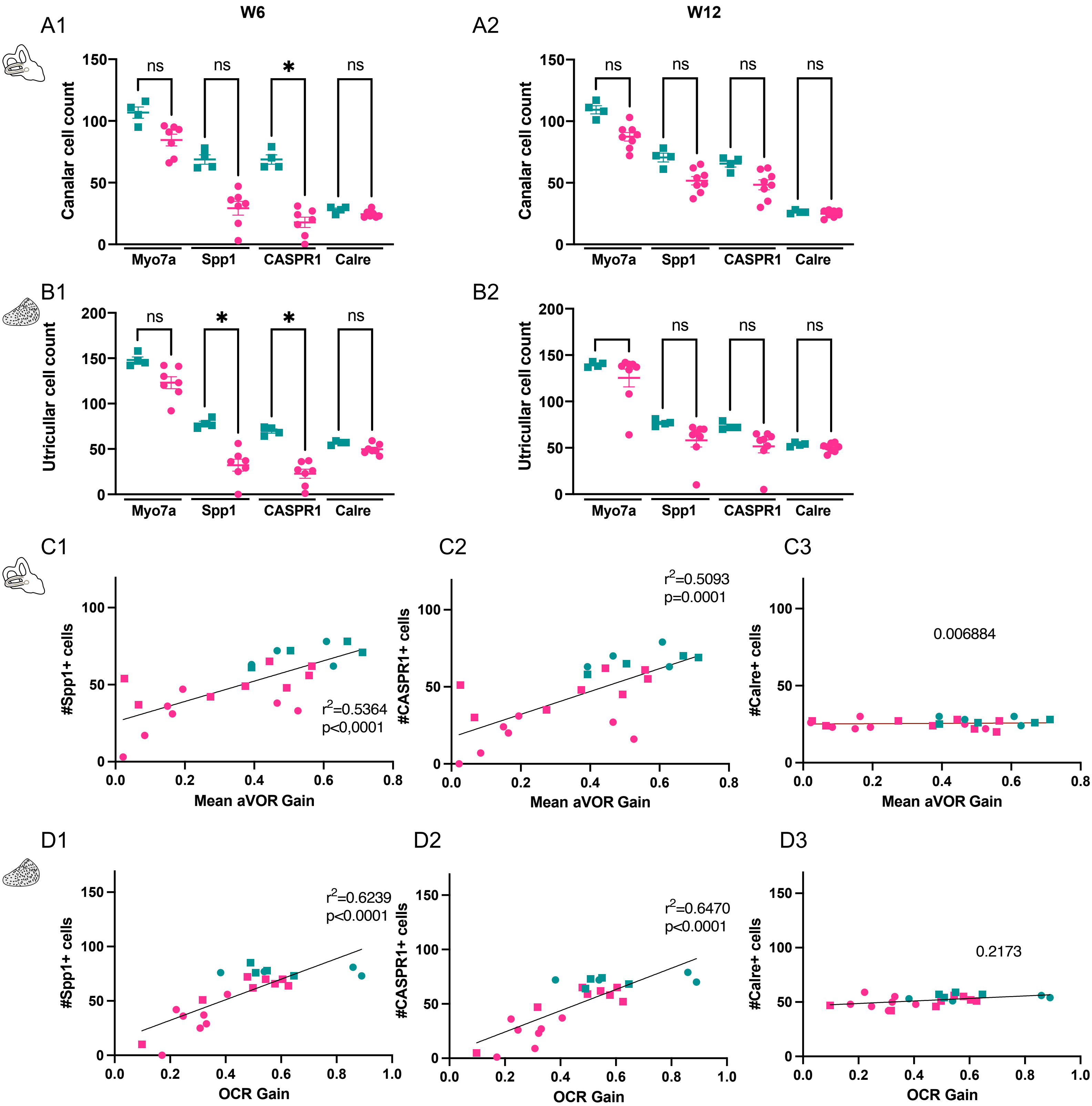


**Figure 3 Supplement: Effects of the IDPN on the number of hair cells in the peripheral regions of the horizontal SCC ampulla and extrastriolar utricule Macula.**

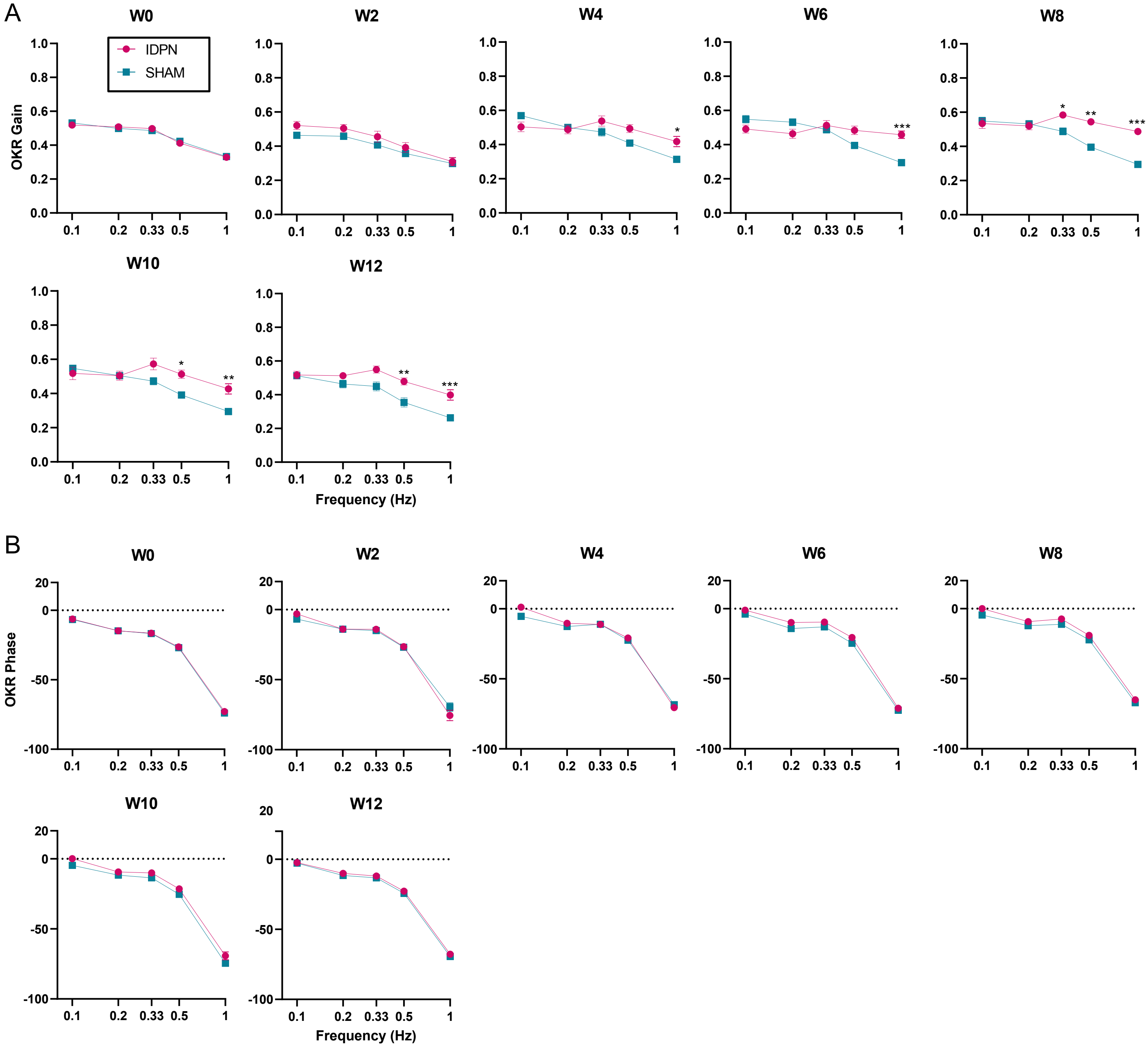


**Figure 4 supplement: Effect of the IDPN on optokinetic reflex amplitude and timing.**
